# Ancient Thioredoxins Evolved to Modern Day Stability-Function Requirement by Altering Native State Ensemble

**DOI:** 10.1101/275982

**Authors:** Tushar Modi, Jonathan Huihui, Kingshuk Ghosh, Sefika Banu Ozkan

## Abstract

Thioredoxins (Thrxs) - small globular proteins that reduce other proteins - are ubiquitous in all forms of life, from archaea to mammals. Although ancestral Thioredoxins share sequential and structural similarity with the modern day (extant) homologs, they exhibit significantly different functional activity and stability. We investigate this puzzle by comparative studies of their (ancient and modern day Thrxs’) native state ensemble, as quantified by the Dynamic Flexibility Index (DFI), a metric for the relative resilience of an amino acid to perturbations in the rest of the protein. Clustering proteins using DFI profiles strongly resembles an alternate classification scheme based on their activity and stability. The DFI profiles of the extant proteins are substantially different around the α3, α4 helices and catalytic regions. Likewise, allosteric coupling of the active site with the rest of the protein is different between ancient and extant Thrxs, possibly explaining the decreased catalytic activity at low pH with evolution. At a global level, we note that the population of low flexibility (called hinges) and high flexibility sites increases with evolution. The heterogeneity (quantified by the variance) in DFI distribution increases with the decrease in the melting temperature typically associated with the evolution of ancient proteins to their modern-day counterparts.

## Introduction

Modern proteins have evolved through small changes from ancient times. Much of this information is encoded in protein classes from different species in the three kingdoms of life (bacteria, archaea, and eukarya). With the advances in phylogenetics and DNA-synthesis techniques, the various ancient genes, including those from the last common ancestors of bacteria, bilaterian animals, and vertebrates, have been resurrected in the laboratories. These studies have provided crucial insights about the environmental adaptations and the evolution of functions [1–8]: (i) ancestral proteins are more robust showing high thermal and chemical stability [9–13], and more interestingly, (ii) protein structures are conserved more than protein sequences throughout the molecular evolution [12–15]. Thus, the current challenge in molecular evolution is to understand the molecular mechanism of how nature alters the function and biophysical properties through amino acid substitutions, while conserving the 3-D structure.

In parallel to the advancements in phylogenetics, recent biophysical studies of proteins have shown that all positions are dynamically linked to each other within a network of interactions, where the strength of each link varies across the protein. This network of interaction leads to intrinsic fluctuations - encoded in the structure and sequence - that govern protein function [4,16– 32]. The obsolete view of the single native structure has been long replaced by “an ensemble of substates” that accurately represent the native state [25]. In order to shed light into the mechanism of evolution and study how evolution shapes the native ensemble, we have developed a method called Dynamic Flexibility Index (DFI) [33]. DFI *is a position specific measure that quantifies* the resilience of a given position (amino acid) to the perturbations occurring at various parts of the protein using linear response theory. Hence, it mimics the multidimensional response when the protein’s conformational space is probed upon interaction with small molecules or other cellular constituents. The DFI is related to a given site’s relative contribution to the conformational entropy of the protein. Because it is a position specific metric, it also allows us to quantify the change in flexibility per position throughout evolution. The DFI identifies flexible and rigid positions within the 3-D interaction network of the protein structure. The low DFI sites are rigid sites (i.e. hinge sites) and are robust to perturbation due to their interaction network within the 3-D structure. However, they efficiently transfer perturbations to rest of the protein chain, similar to joints in a skeleton. Thus, they play critical role in conformational dynamics, and usually corresponds to functionally critical, conserved sites in a protein [33]. On the other hand, high DFI sites are flexible, thus mutations/substitutions on these highly flexible sites are more acceptable. Our earlier work of DFI analysis on over a hundred human proteins has shown that there is a strong positive correlation between DFI and evolutionary rates. This demonstrates that rigid sites are more conserved while highly evolving sites correspond to sites with high flexibility [33]. DFI analysis of evolution of different protein families including GFP proteins [34], beta-lactamase inhibitors [12], and nuclear receptors [35] has shown that alteration of conformational dynamics through allosteric regulations leads to functional changes. Furthermore, our site-specific dynamics-based metric, Dynamic Coupling Index (DCI), reveals that enzymatic function is regulated by dynamically-coupled residues, which form an allosteric communication network with the active sites. Evolution utilizes substitutions on these sites to regulate dynamics of the active sites and binding interfaces [34,36].

Here we analyzed the evolution of Thioredoxin (Thrx). Thioredoxins are versatile small globular protein molecules comprised of about 108 amino acid residues. They belong to a class of oxidoreductase enzymes present in all living organisms from Archaebacteria to Humans. These are labeled as the ultimate *moonlighting* proteins with functions as ubiquitous as being a reducing agent for other proteins in biological reactions [37,38].

Thrxs from Archea to Humans share about 27-69% sequence similarities and a common three-dimensional fold with a central β-sheet core surrounded by four α helices [13,39]. The structure of the reduced and oxidized states of Thrx has been studied extensively over the last few decades. Their structures contain a highly conserved functional site comprising of two neighboring redox active cysteines, Cys-Gly-Pro-Cys (CGPC) [40,41]. The cysteines participate in various redox reactions. The protein is maintained in its oxidized state with its functional site having two thiols with cysteines using another class of reducing agents called Thioredoxin reductases (ThxR) like Nicotinamide adenine dinucleotide phosphate (NADP), Flavin adenine dinucleotide (FAD), etc. [38,40,42]. The reduced state of Thrx facilitates the reduction of the target protein while turning itself into an oxidized state with a disulfide bond between its two functional cysteines. Although, the oxidized state and reduced state of the protein are very similar to each other in their fold (with most of the differences localized around the disulfide active site), chemical denaturation [43] and thermal [44,45] unfolding experiments suggest that Thrxs are more stable in their oxidized state. The mutagenesis analyses of Thrxs have shown that the C-terminal α helix, α4, is known to play a critical role in the folding kinetics of the protein [46]. A shorter helix, α3, has been identified as important for the thermal stability of the fold, having several stabilizing mutations [47]. The central core with β-strand, β5 acts like a bridge for interaction between the two α helices [47] (Figure 1).

**Figure 1.**
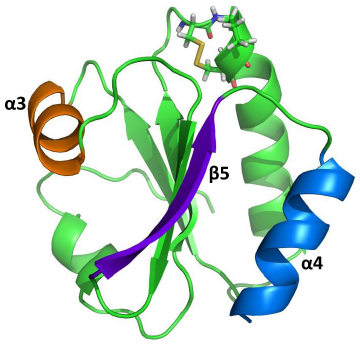
Cartoon representation of AECA Thrx. The active sites, Cys-Gly-Pro-Cys (CGPC) are shown in stick representation, the β-strand, β5, is shown in purple, and the α-helices, α3 in orange and α4 in blue.

Despite a significant difference in the sequences between ancestral and extant enzymes, the ancestral Thioredoxins, which existed almost 4 Billion years ago, display the canonical Thioredoxin fold with only minimal structural changes [3,39,48]. However, during 4 billion years of evolution, the stability of Thrxs have decreased to adapt to the cooling temperatures of the Earth (in mesophilic lineage). Moreover, the catalytic rates of ancestral Thrxs at low pH are different from those of extant ones.

In summary, Thrxs achieved adaptation to a cooler and less acidic Earth by altering their stability and changing their catalytic rates while maintaining the same 3-D fold. In order to provide insights into how variation in sequence alters stability and function while conserving the 3-D structure, we explore how the native state ensemble has evolved throughout evolution of Thrx. To this aim, we performed one-microsecond long Molecular Dynamics simulations for all ancestral and extant Thrxs, and obtained their Dynamic Flexibility Index (DFI) profiles. Comparison of the DFI profiles between ancestral and extant Thrxs on each branch shows a common pattern. The α3 helix, which contributes the most to stability, exhibits enhanced flexibility in modern Thrxs, correlating with the decrease in stability observed in modern enzymes. The increased flexibility of α3 was also associated with increased rigidity in α4 helix along with conserved rigidity of the beta-sheet core. This may have helped to maintain the 3-D fold through the evolution of Thrxs. The clustering of the DFI profiles of all 9 Thrxs and mapping that onto a 2-D landscape of experimentally measured stability and catalytic rates suggests that the native ensemble of Thrxs holds the clue to adaptation at cooler ambient temperature and lower acidic environment.

## Results and Discussion

### The change in DFI profiles provides insight about decrease in melting temperatures during evolution of Thrx

We notice a marked difference between the flexibility profiles of the ancestral and those of the extant Thrxs, particularly the flexibilities of helices α3 and α4. Through mutagenesis analysis it has been shown that disruption of α3 impacts the overall stability of the Thrxs [47]. Interestingly, we observed the enhancement in the flexibility of α3 in EColi (Figure 2).

**Figure 2.**
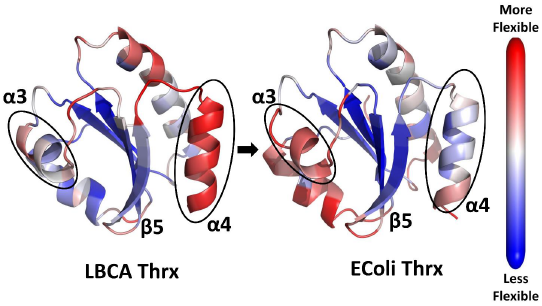
Cartoon representations of ancestral and extant Thioredoxins from bacterial thioredoxins LBCA and EColi Thrx respectively, color coded with their DFI profiles. Red sites are the most flexible and blue site are the least flexible (i.e. rigid). We observe a shift in the flexibility profiles of the helices α3 and α4 through evolution owing to changes in thermal stability. In contrast, the β-sheet core remains rigid throughout evolution, highlighting its importance in mediating the interaction between α3 and α4 helices necessary for the Thrx fold.

DFI profile comparison between LBCA and EColi (Figure 2) shows flexibility-rigidity compensation between the α3 helix and the α4 helix. Specifically, we notice that the α3 helix in EColi has substantially higher DFI values as compared to LBCA, while the opposite is seen in the α4 helix. However, the core region of the β-sheet does not show any noticeable alteration in flexibility between EColi and LBCA besides a slight enhancement in rigidity of β5 β-strand.

These observations can be further correlated with the measured differences in stability due to differences in amino acids in these regions. After performing a sequence alignment between EColi, LBCA, and LPBCA, we note that the ancestral protein LPBCA (close homolog of LBCA) has a critical mutation, P68A (EColi position 68 versus the LPBCA aligned position), in the α3 helix region when compared to EColi. This substitution is typically stabilizing [47]. In the context of two other critical mutations, G74S and K90L, seen in LPBCA, it has been hypothesized that the α3 helix may tilt towards Arginine (R) 89 (another mutation in LPBCA), allowing for a strong charge-dipole interaction [47]. However, EColi Thrx does not allow such favorable interactions due to the different amino acids in these positions. Particularly, 89 is Threonine (T) instead of Arginine (R) in EColi. Thus, the loss of this favorable interaction may explain the lowered stability of EColi Thrx and the enhanced flexibility of the α3 helix as evident from their DFI profiles.

We tested this hypothesis in the context of LBCA. Similar to LPBCA, we note that LBCA also has all the three critical mutations, i.e. P68A, G74S and K90L. Furthermore, after alignment, we see that position 89 has a positively charged amino acid, Lysine, in LBCA.

Motivated by these similarities between LBCA and LPBCA, we further analyzed the relative orientation between the α3 helix dipole and the side chain of Lysine(K) (at the equivalent of position 89) in LBCA. We observed that the relative orientation between the α3 helix dipole and K89 (as seen in sequence alignment) is more rigid in LBCA (Figure SI-1). Whereas, T, at position 89 in EColi, has a fluctuating orientation with the α3 helix (Figure SI-1). This difference indicates that a favorable interaction between the charge and the dipole (of the α3 helix) may be operative in LBCA - similar to LPBCA - and responsible for the higher stability of LBCA and also the rigidity of its α3 helix as compared to EColi Thrx.

Turning our attention to the α4 helix, guided by mutagenesis experiments [47], we again noticed a set of key mutations between EColi and LPBCA responsible for folding stability differences between the two proteins. For example, S95P and Q98A - both occurring near the end of the α4 helix in LPBCA - cause a reduction in stability. It has been hypothesized that both the Serine (S) and Glutamine (Q) in EColi may stabilize the loop connecting the α4 helix and the β5 β-strand by possibly utilizing dipole-dipole interactions. However, upon mutating to Alanine (A) (for Q98A) and Proline (P) (for S95P) in LPBCA these dipolar interactions are lost causing a decrease in stability and consequently high flexibility. A second set of mutations L94Q and F102R (also present in the α4 helix region) in LPBCA have been further implicated to lowering the melting temperature by possibly destabilizing the hydrophobic network between the α4 helix and the β5 β-strand [47]. All these mutations are also present in LBCA, as determined by the sequence alignment, apart from, L94R, present in LBCA instead of L94Q in LPBCA. This suggests that the same physical principles, i.e. loss of charge-dipole or dipole-dipole interaction and/or disruption of the hydrophobic network, may be responsible for the higher flexibility of the α4 helix in LBCA compared to EColi, as observed by DFI analysis.

Similar changes, particularly the compensation for the change in DFI profiles of the α3 and α4 helices, have also been observed in the evolutionary branch of Human Thrxs when the DFI profiles of AECA Thrx and Human Thrx were compared [47] (Figure SI-2). Overall, based on the mutagenesis analysis [47] and DFI comparison between ancestral and extant Thrx suggests a plausible mechanism of how Thrx has evolved to lower stability. While the increased flexibility of the α3 helical region achieved a decrease in stability to adapt to cooler ambient temperatures, the rigidity conservation of the core and increased rigidity in helix α4 ensure conservation of the canonical Thrx fold throughout evolution. The proposed mechanism should be further verified with experimental analysis in the future.

### Dynamic coupling of the active site alters throughout the evolution

Comparison between the catalytic rates of the last ancestral Thrx and their extant variant on each branch in phylogeny shows that the kinetic rates of disulfide bond reduction have decreased during the evolution at pH 5 [3]. Particularly, in the human branch, there is an approximately six-fold decrease in the kinetic rates between its first ancestor, AECA, and the modern-day Human Thrx. Our earlier work on protein evolution shows that nature utilizes distal sites that are dynamically coupled with the active sites to control the active site’s dynamics [49–51]. Thus, we analyzed how the Dynamic Coupling Index (DCI, see methods section for the definition) of the active site changed during Thrx evolution (Figure 3A).

**Figure 3.**
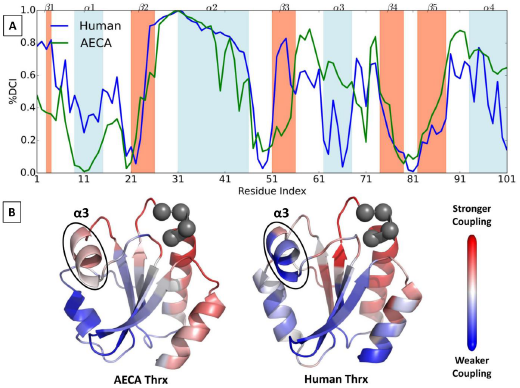
(A) Comparison of the coupling of catalytic sites (CGPC) in ancestral Thrx (AECA) with extant Human Thrx using a percentile ranking of Dynamic Coupling Index (%DCI). We observe a striking difference in the couplings of the α3 region in the respective proteins with their corresponding active sites, suggesting their role in altering catalytic rates. This difference can also be visualized on the cartoon representations (B) of AECA and Human Thrxs where red represents sites highly coupled to the active sites, Cys-Gly-Pro-Cys (CGPC) (grey spheres) and blue as sites with no significant coupling.

Interestingly, we observed that the dynamic coupling of the α3 helix with the active site has decreased drastically between the ancestral and extant Thrxs in the human branch (Figure 3A-B). We noticed the same difference between the ancestral and extant Thrxs in the EColi branch (Figure SI-3). Previous studies have also shown that the α3 helix is part of the substrate binding region and the change in dynamics of this binding region plays a critical role in the catalytic activity of the protein [52]. Thus, the decrease in the allosteric dynamic coupling of the α3 helix region with the catalytic site may be linked with the lowered activity observed in modern day Thrxs. However careful mutagenesis experiment is needed to conclusively prove or disprove this hypothesis.

### The variance in DFI profile distribution correlates with change in melting temperature

DFI analysis of an ancestral versus extant Thrxs in the EColi and Human branches provides insight about how the ‘fine tuning’ of flexibility profiles of some functionally important structural features is utilized during evolution. Comparing the distribution of flexibility of various residue positions in ancestral Thrxs with the extant ones can additionally reveal information about the change in their native landscape through evolution. The distribution of the flexibility of residues in each Thrx protein is obtained by binning the residues according to their DFI scores (Figure 4).

**Figure 4:**
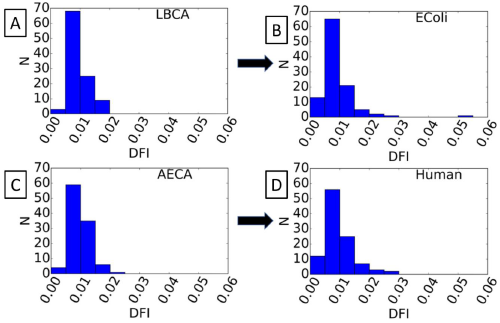
The distributions of DFI profiles in the ancestral and extant Thrx proteins belonging to the ancestor of the Bacterial branch, LBCA (A), evolving to EColi (B), and the ancestor of the Human branch, AECA (C), evolving to extant Human Thrx (D). We observe that through evolution, the residues populate low flexibility and high flexibility regions in the distribution making it wider. The fraction of residues populating high flexibility (DFI greater than 0.02) regions increased in EColi from 0% ∼4% and in Human from 1% to 4%

Interestingly, the distribution of DFI profile for the 4Gyr old Thrx (LBCA) differs from that of extant Thrx (EColi) (Figure 4A-B). The high flexibility tail region of the distribution gets more populated in modern Thrxs. Likewise, the probability density at low DFI has also increased in Thrx of modern organisms, representing a gain of high flexibility and low flexibility regions during evolution. This behavior was observed not only in the Bacterial but also in the Human branch of Thrx’s evolutionary tree (Figure 4C-D). The redistribution of flexible sites in Thrx structures help proteins ‘fine tune’ their activity in accordance with the functional requirement.

This characteristic pattern of increasing ‘width’ of the distributions with evolution is further supported by the high correlation between the variance of the DFI distributions of Thrx proteins and their corresponding melting temperatures (correlation, R=-0.86 and p=3.2×10^−3^) (Figure 5A). In addition, since the decreasing melting temperature correlates with the evolution time, we also notice significant correlation between the evolution time of Thrx proteins and their variance in DFI distributions (correlation, R=0.77 and p=1.6×10^−2^) (Figure 5B)

**Figure 5:**
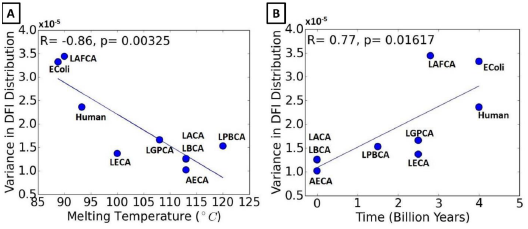
(A) The variance in distribution of the flexibility profiles in ancestral and modern day Thioredoxin proteins are observed to correlate strongly with their melting temperatures [8], R=- 0.86, indicating the implication of flexibility profiles in their thermostability. (B) The variance in DFI distributions is observed to increase with evolution [8] and is strongly correlated with the time of evolution of the proteins, R=0.77.

Overall, this result is in agreement with our previous results [12,34] that evolution shapes the conformational landscape of the native state. We observed two major changes in DFI distributions consistent in all branches. First, the DFI profiles change as the sequences evolve. Second, the probability distributions at the beginning (ancestral sequences) are more compact, having a higher probability towards mid to high DFI values. This type of distribution ensures evolvability through mutation of various flexible positions in the protein since flexible sites typically correspond to highly evolving residues [53–55]. On the other hand, as we get closer to the modern enzymes, the distribution widens. There is an increase in the probability of the low DFI range along with a longer tail of high DFI values. In other words, a well-distributed set of very rigid and very flexible sites could be an evolutionary mechanism to adapt to low temperature and/or adjust to functional need.

### Dynamic Flexibility Index (DFI) captures the functional evolution in thioredoxin

To test how the DFI profiles of nine Thrxs captures the change in function throughout evolution, we clustered their DFI profiles using PCA analysis (See methods). The Thrxs that clustered together in accordance with the similarities in their flexibility profiles (Figure 6A) also exhibit similar rate constants for disulfide bond reduction obtained from single molecule experiments [3]. For example, ancestral Thrxs LACA and AECA, belonging to the Archaea branch, have very high rate constants. Consistently, DFI clustering arranges them in the same group. On the other hand, LBCA Thrx, from the Bacterial branch, which evolved around the same period as AECA (around 4 Billion year ago), is in a different cluster with LPBCA and LGPCA. LBCA, LPBCA, and LGPCA all have much lower rates for disulfide bond reduction than those of LACA and AECA. Interestingly, Human, LAFCA, and LECA Thrx, all grouped together in a common cluster, share similar kinetic rates of disulfide bond reduction. We also notice EColi, having the lowest reduction rate of all other Thrx, is clustered in a separate group (Figure 6A).

**Figure 6:**
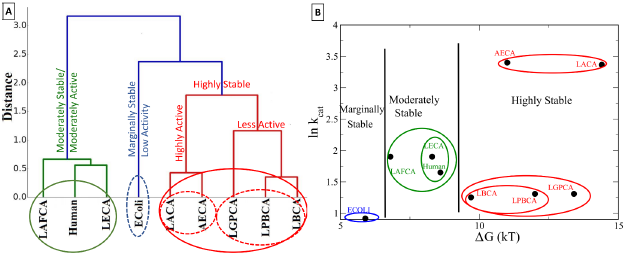
Clustering of ancestral and extant Thioredoxins based on their DFI profiles (A) and based on their experimental rate constants of disulfide bond reduction [56] and change in Free energy of folding ΔG_folding_ [56] (B). The two clustering criteria give similar results as they segregate the nine Thioredoxin proteins based on their activities and thermal stability.

While the above argument clearly explains subclasses, the clustering also raises question about merging different subgroups to bigger groups. For example, how do we rationalize that LACA-AECA dyad is grouped in the same class (red) with the triad LGPCA, LPBCA-LBCA in spite of the two subclasses having widely different catalytic rates? Interestingly, we notice LACA, AECA, LGPCA, LPBCA, and LBCA are all highly stable compared to the triad LAFCA, Human, and LECA. Thus, DFI based classification can be best understood by both catalytic rates (kcat, M^-1^s^-1^) [3] and free energy of folding [56]. It is interesting to note that although DFI solely utilizes the native state ensemble, it can successfully sculpt the landscape in these two coordinates (Figure 6B). Based on stability and catalytic activity, Thrxs can be grouped into three broad categories: (i) highly stable agents (AECA and LACA with high activity, LBCA, LPBCA, and LGPCA with low activity), (ii) moderately-stable agents (Human, LAFCA, and LECA) with moderate catalytic activity and (iii) marginally stable, low catalytic activity (EColi) (Figure 6A, B).

In summary, clustering solely based on DFI profiles successfully captures the clustering based on catalytic rates, and stability. This suggests that evolution shaped the native state ensemble of Thrxs to adapt and function at cooler temperatures and lower acidic ambient conditions. It is also in agreement with our previous analysis on protein evolution [12,50], highlighting that evolution exploits native state conformational dynamics to alter function.

## Conclusion

Despite the significant structural similarities between ancestral and extant Thioredoxins, they evolved towards lower stability and kinetic turnover rates. In order to gain insights to the underlying molecular mechanism, we explored how changes in the native state ensemble might have impacted the evolution of Thrx proteins. To this aim, we compared the difference in their Dynamic Flexibility Index (DFI) profiles. The enhanced flexibility of the α3 helix in the extant proteins, compared to their ancestral counterparts, is compensated by the decrease in flexibility of the α4 helix and may be responsible for lowering their stability to adapt to cooler ambient temperatures, while keeping the fold conserved. We further notice that the dynamic coupling of important positions with the catalytic site has changed during evolution. Particularly, the decrease in allosteric dynamic coupling of the α3 helical region - critical in substrate binding - with the catalytic site in extant Thrxs may be associated with the decrease in catalytic activity at lower acidic conditions.

Comparison of the distribution of the flexibility of residues between ancestral and extant proteins revealed that the population density of high and low flexibility sites increases as they evolve. These common features observed in evolution suggest a “*fine tuning*” of their native ensemble to adjust to ambient conditions in accordance with the evolution in their function. The high correlation between the variance of flexibility distribution of proteins and their melting temperature quantitatively supports this hypothesis.

In addition, clustering these proteins based on their flexibility profiles closely matches grouping using their kinetic rates of disulfide bond reduction and thermal stability. These observations, in agreement with our previous results, highlight that nature utilizes native state conformational dynamics to adapt and evolve.

## Method

### Molecular Dynamics simulation protocol

All starting structures were taken from the PDB using the respective ID’s for the proteins LBCA (4BA7), AECA (3ZIV), LAFCA (2YPM), LECA (2YOI), LACA (2YNX), LGPCA (2YN1), LPBCA (2YJ7), EColi (2TRX), and Human (1ERU) Thrx [57–59]. Next the H++ web server was used to predict the protonation state of the histidine side chains [60–62]. The refined structure was then loaded into TLEAP using the ff14SB force field [63]. The disulfide bond was kept in the oxidized state. Protein hydrogens were then added and a 9.0 Angstrom cubic box of TIP3P surrounding water atoms was added, followed by neutralizing ions [64,65]. Systems were then energy minimized using the SANDER module of AMBER 14 [66–68]. The first cycle of minimization reduced the energy and steric clashes of the solvent with the protein restricted using harmonic restraints. The second cycle of minimization was then performed without the harmonic restraints so the entire solution could adjust to the local minimum.

Heating, density equilibration, and production were all then run using the GPU-accelerated PMEMD module of AMBER 14 [68]. These simulations were performed with periodic boundary conditions and constrained the bond lengths of all covalent hydrogen bonds using SHAKE [67]. Direct-sum, non-bonded interactions were cutoff at 9.0 Angstroms, and long-range electrostatics were calculated using the Particle Mesh Ewald method [69–71]. The heating phase ran over 100 picoseconds from 0 to 300K. The density of the system was then allowed to equilibrate over 5 nanoseconds at constant temperature and pressure. All production simulations were run using the Langevin thermostat and Berendsen barostat to keep temperature and pressure, respectively, constant. A time step of 2 femtoseconds was used and structural conformations were saved every 10 picoseconds. All simulations were allowed to progress to 1 microsecond of total simulation time deemed minimal required simulation time for convergence based on our earlier study [72].

### Dynamic Flexibility Index (DFI)

DFI measures the amount of resilience a residue experiences in response to perturbations in the rest of the protein. The dynamic response profile of the protein is explored using Perturbation Response Scanning technique (PRS) [73,74]. The original approach is based on Elastic Network Model (ENM) in which the nodes represent Cα atoms [54,75] and the pairwise potential between each atom is given by the potential of a harmonic spring. A small perturbation in the form of random Brownian kick is applied sequentially to each Cα atom in the elastic network. As a first order approximation, this perturbation mimics the forces exerted by an approaching protein or a ligand in a crowded cellular environment. The perturbations on a single residue result in a cascade of perturbations to all other atoms in the network, inducing a global response. The fluctuation response profile of the positions upon perturbation of a single residue is obtained using linear response theory and given by the equation:

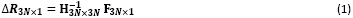

Where, **H** is the hessian matrix 3Nx3N matrix composed of the second order derivatives of the harmonic potential with respect to the components of the position vectors for the chain of length N, giving the position co-variance of the residue pairs in equilibrium, **F** is the external unit force vector applied at N residues in the protein and **ΔR** is the response of the force. The force is applied in all the directions at each residue and the magnitude of response profile is averaged to give an isotropic measure of response.

However, a disadvantage of the ENM-based PRS model is that the coarse-grained network makes it insensitive to changes arising from the biochemical properties of amino acids. Therefore, in order to compare the family of Thrxs, having similar back-bone structures, we replace the inverse of Hessian with the covariance matrix obtained from molecular dynamic simulations (discussed in the previous section).

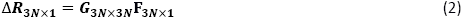

Here, **G** is the covariance matrix containing the dynamic properties of the system. The covariance matrix contains the data for long range interactions, solvation effects, and biochemical specificities of all type of interactions. The DFI is calculated by computing the fluctuation response of all the residues in a protein (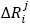, *i* = 1,.. *N*) by applying a unit force (isotropic) as perturbation at a specific position (*i.e.* site j) using equation 2. We repeat this single residue perturbation for all residues (i.e. 1≤j≤N;) in the protein. Consequently, the Perturbation Matrix, A is constructed,

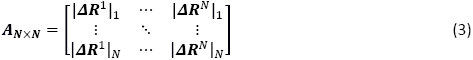

Where,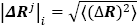 is the magnitude of fluctuation response at site ‘i’ due to the perturbations at site ‘j‘. A sum of a given row of perturbation matrix gives the net average displacement of the residue from its equilibrium position when all the residues are perturbed by an isotropic unit force one at a time. The DFI score of a position ‘i’ is defined as the net response of that position normalized with the net displacement of the whole protein when all the residues are perturbed, i.e.

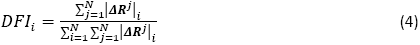

Thus, a higher DFI score of a residue position ‘i’ implies a more flexible site and a low score implies a rigid site with lower response to perturbations in the protein.

Recently, we have extended this method to identify dynamic coupling between any given residue and functionally important residues by introducing a new metric called the dynamic coupling index (DCI). DCI metric can identify sites that are distal to functional sites but impact active site dynamics through dynamic allosteric coupling [49,50,76]. This type of allosteric coupling is important; sites with strong dynamic allosteric coupling to functionally critical residues (Dynamic Allosteric Residue Coupling (DARC) spots), regardless of separation distance, contribute to the function as well. Thus, a mutation at such a site can disrupt the allosteric dynamic coupling or regulation, leading to functional degradation. As defined, the DCI is the ratio of the sum of the mean square fluctuation response of the residue ‘i’ upon functional site perturbations (i.e., catalytic residues) to the response of residue ‘i’ upon perturbations on all residues. DCI enables us to identify DARC spot residues, which are more sensitive to perturbations exerted on residues critical for function. This index can be utilized to identify the residues involved in allosteric regulation. It is expressed as:

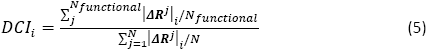

here |Δ**R**^j^|_i_ is the response fluctuation profile of residue ‘i’ upon perturbation of residue ‘j‘. The numerator is the average mean square fluctuation response obtained over the perturbation of the functionally critical residues N_functional_and the denominator is the average mean square fluctuation response over all residues. Similar to DFI, the DCI profiles can also be computed using the covariance matrix obtained from Molecular Dynamics simulations or the inverse Hessian of the elastic network model. In this study, we used the covariance matrices.

### Clustering the DFI profiles of Thioredoxin proteins

We clustered the DFI profiles of different extant and ancestral and Thioredoxins by comparing their percentile rankings. In order to compare the flexibility profiles, the proteins are aligned according to their multiple sequence alignment and are concatenated into a data matrix **X**. Singular Value Decomposition (SVD) is a statistical procedure to factorize the data into the orthonormal basis which represent the vector space of data. It is similar to Principal Component Analysis which could be used to understand the structure of data or to increase the signal to noise ratio in data by eliminating the redundant dimensions and mapping it on a lower dimensional space. Clustering by SVD acts as an effective noise filter by isolating the highest variances among data points in the top principal vectors. Consequently, the remaining insignificant singular vectors can be omitted from the reconstruction.

The DFI profiles of all proteins are merged into a matrix, **X**, of dimensions (**m** × **n**). Here **m** is the number of datasets (proteins) we are clustering together, each having **n** number of attributes (number of residues). On performing SVD, **X** is decomposed as follows:

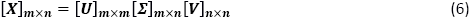

Here, **U** and **V** are unitary matrices with orthonormal columns and are called left singular vectors and right singular vectors respectively, and Σis a diagonal matrix with diagonal elements known as Singular values of **X**.

The singular values of **X**, by convention, are arranged in a decreasing order of their magnitude, σ = {σ_i_} represent the variances in the corresponding left and right singular vectors. The set of highest singular values representing the largest variance in the orthonormal singular vectors can be interpreted to show the characteristics in the data **X** and the right singular vectors create orthonormal basis which spans the vector space representing the data. The left singular vectors contain weights indicating the significance of each attribute in the dataset as: 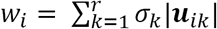 Using these features of the decomposed singular vectors, we can create another matrix, X* using only the highest ‘r’ singular values which can mimic the basic characteristics of the original dataset. Thus, X* can be represented as:

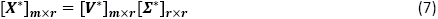

Here, Σ* contains only largest ***r*** singular values and V*contains the corresponding right singular vectors. The data is now clustered hierarchically based on the pairwise distance between different proteins in the reconstructed DFI data with reduced dimensions.

For a pair of datasets (or between flexibility profiles of any two proteins) j_1_ and j_2_, the distance between them in original set of data was given by:

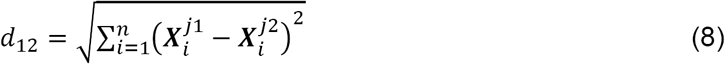

Which in reduced dimensions can be calculated as:

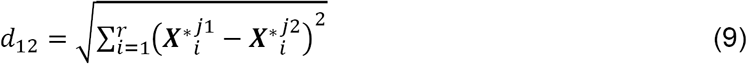

These pairwise distances are used as the parameters for clustering the flexibility profiles of Thioredoxin. We calculated the multiple sequence alignment of ancestral and modern Thrx. The DFI profiles are aligned with respect to LBCA taking into consideration the gaps in sequences of other Thrx proteins, and the data is clubbed into a dataset matrix **X**. Three largest singular values are used for reconstruction of data and clustering. The pairwise distance between each protein using the equation above is used for clustering them hierarchically.

A bottom up approach is used for the hierarchical clustering where initially each protein is assigned its own cluster and then in successive iteration, closest clusters are merged together into a common cluster. In this approach, the distance between clusters is defined by the average pairwise distance between their components (Average Linkage clustering [77]). In the end, the clusters are represented hierarchically using a dendrogram, where the vertical axis denotes the Euclidian distance between various clusters and among their sub-clusters.

## Acknowledgment

Support from NSF-MCB Award 1715591 and Scialog Fellow Award by RCSA and the Gordon & Betty Moore Foundation is gratefully acknowledged by SBO. KG acknowledges support from NSF (award number 1149992) and Research Corporation for Science Advancement. We thank Lucas Sawle for the help with the all-atom molecular dynamics simulation.

## Author Contributions

SBO and KG. conceived and designed the study. TM and JH performed and analyzed simulations, and generated the results. TM, JH, KG and SBO wrote the manuscript

## Bibliography

1. Carroll SM, Bridgham JT, Thornton JW. Evolution of Hormone Signaling in Elasmobranchs by Exploitation of Promiscuous Receptors. Mol Biol Evol. 2008 Dec;25(12):2643–52.

2. Ortlund EA, Bridgham JT, Redinbo MR, Thornton JW. Crystal structure of an ancient protein. Science. 2007 Sep 14;317(5844):1544–8.

3. Perez-Jimenez R, Inglés-Prieto A, Zhao Z-M, Sanchez-Romero I, Alegre-Cebollada J, Kosuri P, Garcia-Manyes S, Kappock TJ, Tanokura M, Holmgren A, Sanchez-Ruiz JM, Gaucher EA, Fernandez JM. Single-molecule paleoenzymology probes the chemistry of resurrected enzymes. Nat Struct Mol Biol. 2011 Apr 3;18(5):nsmb.2020.

4. Wilson C, Agafonov RV, Hoemberger M, Kutter S, Zorba A, Halpin J, Buosi V, Otten R, Waterman D, Theobald DL, Kern D. Kinase dynamics. Using ancient protein kinases to unravel a modern cancer drug’s mechanism. Science. 2015 Feb 20;347(6224):882–6.

5. Bar-Rogovsky H, Hugenmatter A, Tawfik DS. The evolutionary origins of detoxifying enzymes: the mammalian serum paraoxonases (PONs) relate to bacterial homoserine lactonases. J Biol Chem. 2013 Aug 16;288(33):23914–27.

6. Smith SD, Wang S, Rausher MD. Functional evolution of an anthocyanin pathway enzyme during a flower color transition. Mol Biol Evol. 2013 Mar;30(3):602–12.

7. Boucher JI, Jacobowitz JR, Beckett BC, Classen S, Theobald DL. An atomic-resolution view of neofunctionalization in the evolution of apicomplexan lactate dehydrogenases. eLife. 2014 Jun 25;3.

8. Ingles-Prieto A, Ibarra-Molero B, Delgado-Delgado A, Perez-Jimenez R, Fernandez JM, Gaucher EA, Sanchez-Ruiz JM, Gavira JA. Conservation of protein structure over four billion years. Struct Lond Engl 1993. 2013 Sep 3;21(9):1690–7.

9. Trudeau DL, Kaltenbach M, Tawfik DS. On the Potential Origins of the High Stability of Reconstructed Ancestral Proteins. Mol Biol Evol. 2016 Oct 1;33(10):2633–41.

10. Akanuma S, Nakajima Y, Yokobori S, Kimura M, Nemoto N, Mase T, Miyazono K, Tanokura M, Yamagishi A. Experimental evidence for the thermophilicity of ancestral life. Proc Natl Acad Sci U S A. 2013 Jul 2;110(27):11067–72.

11. Hart KM, Harms MJ, Schmidt BH, Elya C, Thornton JW, Marqusee S. Thermodynamic system drift in protein evolution. PLoS Biol. 2014 Nov;12(11):e1001994.

12. Zou T, Risso VA, Gavira JA, Sanchez-Ruiz JM, Ozkan SB. Evolution of Conformational Dynamics Determines the Conversion of a Promiscuous Generalist into a Specialist Enzyme. Mol Biol Evol. 2015 Jan 1;32(1):132–43.

13. Risso VA, Manssour-Triedo F, Delgado-Delgado A, Arco R, Barroso-delJesus A, Ingles-Prieto A, Godoy-Ruiz R, Gavira JA, Gaucher EA, Ibarra-Molero B, Sanchez-Ruiz JM. Mutational studies on resurrected ancestral proteins reveal conservation of site-specific amino acid preferences throughout evolutionary history. Mol Biol Evol. 2015 Feb;32(2):440–55.

14. Bridgham JT, Ortlund EA, Thornton JW. An epistatic ratchet constrains the direction of glucocorticoid receptor evolution. Nature. 2009 Sep 24;461(7263):515–9.

15. Choi I-G, Kim S-H. Evolution of protein structural classes and protein sequence families. Proc Natl Acad Sci. 2006 Sep 19;103(38):14056–61.

16. McLeish TCB, Cann MJ, Rodgers TL. Dynamic Transmission of Protein Allostery without Structural Change: Spatial Pathways or Global Modes? Biophys J. 2015 Sep 15;109(6):1240–50.

17. Bahar I, Lezon TR, Yang L-W, Eyal E. Global dynamics of proteins: bridging between structure and function. Annu Rev Biophys. 2010;39:23–42.

18. Bahar I, Chennubhotla C, Tobi D. Intrinsic Enzyme Dynamics in the Unbound State and Relation to Allosteric Regulation. Curr Opin Struct Biol. 2007 Dec;17(6):633–40.

19. Chennubhotla C, Yang Z, Bahar I. Coupling between global dynamics and signal transduction pathways: a mechanism of allostery for chaperonin GroEL. Mol Biosyst. 2008 Mar 18;4(4):287–92.

20. Tobi D, Bahar I. Structural changes involved in protein binding correlate with intrinsic motions of proteins in the unbound state. Proc Natl Acad Sci U S A. 2005 Dec 27;102(52):18908–13.

21. Boehr DD, Dyson HJ, Wright PE. An NMR Perspective on Enzyme Dynamics. Chem Rev. 2006Aug 1;106(8):3055–79.

22. Mazal H, Aviram H, Riven I, Haran G. Effect of ligand binding on a protein with a complex folding landscape. Phys Chem Chem Phys [Internet]. 2017 Jun 26 [cited 2017 Nov 1]; Available from: http://pubs.rsc.org/en/content/articlelanding/2018/cp/c7cp03327c

23. Kar G, Keskin O, Gursoy A, Nussinov R. Allostery and population shift in drug discovery. Curr Opin Pharmacol. 2010 Dec;10(6):715–22.

24. Tsai C-J, Nussinov R. A Unified View of “How Allostery Works.” PLOS Comput Biol. 2014;10(2):1–12.

25. Tokuriki N, Tawfik DS. Protein dynamism and evolvability. Science. 2009 Apr 10;324(5924):203–7.

26. Henzler-Wildman KA, Lei M, Thai V, Kerns SJ, Karplus M, Kern D. A hierarchy of timescales in protein dynamics is linked to enzyme catalysis. Nature. 2007 Dec 6;450(7171):913–6.

27. Zheng W, Brooks BR, Thirumalai D. Low-frequency normal modes that describe allosteric transitions in biological nanomachines are robust to sequence variations. Proc Natl Acad Sci U S A. 2006 May 16;103(20):7664–9.

28. Dima RI, Thirumalai D. Determination of network of residues that regulate allostery in protein families using sequence analysis. Protein Sci Publ Protein Soc. 2006 Feb;15(2):258–68.

29. Liu T, Whitten ST, Hilser VJ. Functional residues serve a dominant role in mediating the cooperativity of the protein ensemble. Proc Natl Acad Sci U S A. 2007 Mar 13;104(11):4347–52.

30. Gruber R, Horovitz A. Allosteric Mechanisms in Chaperonin Machines. Chem Rev. 2016 Jun 8;116(11):6588–606.

31. Buchenberg S, Sittel F, Stock G. Time-resolved observation of protein allosteric communication. Proc Natl Acad Sci. 2017 Aug 15;114(33):E6804–11.

32. Sawle L, Huihui J, Ghosh K. All-Atom Simulations Reveal Protein Charge Decoration in the Folded and Unfolded Ensemble Is Key in Thermophilic Adaptation. J Chem Theory Comput. 2017 Oct 10;13(10):5065–75.

33. Nevin Gerek Z, Kumar S, Banu Ozkan S. Structural dynamics flexibility informs function and evolution at a proteome scale. Evol Appl. 2013 Apr;6(3):423–33.

34. Kim H, Zou T, Modi C, Dörner K, Grunkemeyer TJ, Chen L, Fromme R, Matz MV, Ozkan SB, Wachter RM. A hinge migration mechanism unlocks the evolution of green-to-red photoconversion in GFP-like proteins. Struct Lond Engl 1993. 2015 Jan 6;23(1):34–43.

35. Glembo TJ, Farrell DW, Gerek ZN, Thorpe MF, Ozkan SB. Collective Dynamics Differentiates Functional Divergence in Protein Evolution. PLOS Comput Biol. 2012 Mar 29;8(3):e1002428.

36. Kumar A, Butler BM, Kumar S, Ozkan SB. Integration of structural dynamics and molecular evolution via protein interaction networks: a new era in genomic medicine. Curr Opin Struct Biol. 2015 Dec 1;35(Supplement C):135–42.

37. Holmgren A. Thioredoxin. Annu Rev Biochem. 1985;54(1):237–71.

38. Mustacich D, Powis G. Thioredoxin reductase. Biochem J. 2000 Feb 15;346(Pt 1):1–8.

39. Romero-Romero ML, Risso VA, Martinez-Rodriguez S, Ibarra-Molero B, Sanchez-Ruiz JM. Engineering ancestral protein hyperstability. Biochem J. 2016 Oct 15;473(20):3611–20.

40. Eklund H, Gleason FK, Holmgren A. Structural and functional relations among thioredoxins of different species. Proteins. 1991;11(1):13–28.

41. Weichsel A, Gasdaska JR, Powis G, Montfort WR. Crystal structures of reduced, oxidized, and mutated human thioredoxins: evidence for a regulatory homodimer. Structure. 1996 Jun 1;4(6):735–51.

42. Arnér ESJ, Holmgren A. Physiological functions of thioredoxin and thioredoxin reductase. Eur J Biochem. 2000 Oct 1;267(20):6102–9.

43. Chakrabarti A, Srivastava S, Swaminathan CP, Surolia A, Varadarajan R. Thermodynamics of replacing an α-helical pro residue in the p40S mutant of Escherichia-coli thioredoxin. Protein Sci. 1999;8(11):2455–9.

44. Ladbury JE, Kishore N, Hellinga HW, Wynn R, Sturtevant JM. Thermodynamic effects of reduction of the active-site disulfide of Escherichia coli thioredoxin explored by differential scanning calorimetry. Biochemistry (Mosc). 1994 Mar 29;33(12):3688–92.

45. Godoy-Ruiz R, Perez-Jimenez R, Ibarra-Molero B, Sanchez-Ruiz JM. Relation Between Protein Stability, Evolution and Structure, as Probed by Carboxylic Acid Mutations. J Mol Biol. 2004 Feb 13;336(2):313–8.

46. Vazquez DS, Sánchez IE, Garrote A, Sica MP, Santos J. The E. coli thioredoxin folding mechanism: the key role of the C-terminal helix. Biochim Biophys Acta. 2015 Feb;1854(2):127–37.

47. Cabrera ÁC, A. Sánchez-Murcia P, Gago F. Making sense of the past: hyperstability of ancestral thioredoxins explained by free energy simulations. Phys Chem Chem Phys. 2017;19(34):23239–46.

48. Risso VA, Gavira JA, Gaucher EA, Sanchez-Ruiz JM. Phenotypic comparisons of consensus variants versus laboratory resurrections of Precambrian proteins. Proteins. 2014 Jun;82(6):887–96.

49. Larrimore KE, Kazan IC, Kannan L, Kendle RP, Jamal T, Barcus M, Bolia A, Brimijoin S, Zhan C-G, Ozkan SB, Mor TS. Plant-expressed cocaine hydrolase variants of butyrylcholinesterase exhibit altered allosteric effects of cholinesterase activity and increased inhibitor sensitivity. Sci Rep [Internet]. 2017 Sep 5;7. Available from: https://www.ncbi.nlm.nih.gov/pmc/articles/PMC5585256/

50. Kumar A, Glembo TJ, Ozkan SB. The Role of Conformational Dynamics and Allostery in the Disease Development of Human Ferritin. Biophys J. 2015 Sep 15;109(6):1273–81.

51. Butler BM, Kumar A, Gerek ZN, Sanderford M, Kumar S, Ozkan SB. Dynamic coupling of residues in proteins explains enigmatic pathogenic missense variation in humans. Rev.

52. Perez-Jimenez R, Li J, Kosuri P, Sanchez-Romero I, Wiita AP, Rodriguez-Larrea D, Chueca A, Holmgren A, Miranda-Vizuete A, Becker K, Cho S-H, Beckwith J, Gelhaye E, Jacquot JP, Gaucher EA, Sanchez-Ruiz JM, Berne BJ, Fernandez JM. Diversity of chemical mechanisms in thioredoxin catalysis revealed by single-molecule force spectroscopy. Nat Struct Mol Biol. 2009 Aug;16(8):890.

53. Gerek ZN, Ozkan SB. A flexible docking scheme to explore the binding selectivity of PDZ domains. Protein Sci Publ Protein Soc. 2010 May;19(5):914–28.

54. Gerek ZN, Ozkan SB. Change in allosteric network affects binding affinities of PDZ domains: analysis through perturbation response scanning. PLoS Comput Biol. 2011 Oct;7(10):e1002154.

55. Gerek ZN, Keskin O, Ozkan SB. Identification of specificity and promiscuity of PDZ domain interactions through their dynamic behavior. Proteins. 2009 Dec;77(4):796–811.

56. Tzul FO, Vasilchuk D, Makhatadze GI. Evidence for the principle of minimal frustration in the evolution of protein folding landscapes. Proc Natl Acad Sci. 2017 Feb 28;114(9):E1627–32.

57. Risso VA, Gavira JA, Mejia-Carmona DF, Gaucher EA, Sanchez-Ruiz JM. Hyperstability and substrate promiscuity in laboratory resurrections of Precambrian β-lactamases. J Am Chem Soc. 2013 Feb 27;135(8):2899–902.

58. Katti SK, LeMaster DM, Eklund H. Crystal structure of thioredoxin from Escherichia coli at 1.68 A resolution. J Mol Biol. 1990 Mar 5;212(1):167–84.

59. Capitani G, Markovic-Housley Z, Jansonius JN, Val GD, Morris M, Schürmann P, Biochimie L. Crystal Structures of Thioredoxins f and m from Spinach Chloroplasts. In: Photosynthesis: Mechanisms and Effects [Internet]. Springer, Dordrecht; 1998 [cited 2017 Oct 1]. p. 1939–42. Available from: https://link.springer.com/chapter/10.1007/978-94-011-3953-3_452

60. Anandakrishnan R, Aguilar B, Onufriev AV. H++ 3.0: automating pK prediction and the preparation of biomolecular structures for atomistic molecular modeling and simulations. Nucleic Acids Res. 2012 Jul;40(Web Server issue):W537–41.

61. Myers J, Grothaus G, Narayanan S, Onufriev A. A simple clustering algorithm can be accurate enough for use in calculations of pKs in macromolecules. Proteins Struct Funct Bioinforma. 2006 Jun 1;63(4):928–38.

62. Gordon JC, Myers JB, Folta T, Shoja V, Heath LS, Onufriev A. H++: a server for estimating pKas and adding missing hydrogens to macromolecules. Nucleic Acids Res. 2005 Jul 1;33(Web Server issue):W368–371.

63. Maier JA, Martinez C, Kasavajhala K, Wickstrom L, Hauser KE, Simmerling C. ff14SB: Improving the Accuracy of Protein Side Chain and Backbone Parameters from ff99SB. J Chem Theory Comput. 2015 Aug 11;11(8):3696–713.

64. Jorgensen WL, Chandrasekhar J, Madura JD, Impey RW, Klein ML. Comparison of simple potential functions for simulating liquid water. J Chem Phys. 1983 Jul 15;79(2):926–35.

65. Neria E, Fischer S, Karplus M. Simulation of activation free energies in molecular systems. J Chem Phys. 1996 Aug 1;105(5):1902–21.

66. Case D, Babin V, Berryman J, Betz R, Cai Q, Cerutti D, Cheatham T, Darden T, Duke R, Gohlke H, Goetz A, Gusarov S, Homeyer N, Janowski P, Kaus J, Kolossváry I, Kovalenko A, Lee T, LeGrand S, Luchko T, Luo R, Madej B, Merz K, Paesani F, Roe D, Roitberg A, Sagui C, Salomon-Ferrer R, Seabra G, Simmerling C, Smith W, Swails J, Walker, Wang J, Wolf R, Wu X, Kollman P. {Amber 14}. 2014.

67. Pearlman DA, Case DA, Caldwell JW, Ross WS, Cheatham TE, DeBolt S, Ferguson D, Seibel G, Kollman P. AMBER, a package of computer programs for applying molecular mechanics, normal mode analysis, molecular dynamics and free energy calculations to simulate the structural and energetic properties of molecules. Comput Phys Commun. 1995 Sep 2;91(1–3):1–41.

68. Salomon-Ferrer R, Götz AW, Poole D, Le Grand S, Walker RC. Routine Microsecond Molecular Dynamics Simulations with AMBER on GPUs. 2. Explicit Solvent Particle Mesh Ewald. J Chem Theory Comput. 2013 Sep 10;9(9):3878–88.

69. Darden T, York D, Pedersen L. Particle mesh Ewald: An N⋅log(N) method for Ewald sums in large systems. J Chem Phys. 1993 Jun 15;98(12):10089–92.

70. Essmann U, Perera L, Berkowitz ML, Darden T, Lee H, Pedersen LG. A smooth particle mesh Ewald method. J Chem Phys. 1995 Nov 15;103(19):8577–93.

71. Hockney RW, Eastwood JW. Computer Simulation Using Particles. Bristol, PA, USA: Taylor & Francis, Inc.; 1988.

72. Sawle L, Ghosh K. Convergence of Molecular Dynamics Simulation of Protein Native States: Feasibility vs Self-Consistency Dilemma. J Chem Theory Comput. 2016 Feb 9;12(2):861–9.

73. Bolia A, Gerek ZN, Keskin O, Banu Ozkan S, Dev KK. The binding affinities of proteins interacting with the PDZ domain of PICK1. Proteins. 2012 May;80(5):1393–408.

74. Bolia A, Ozkan SB. Adaptive BP-Dock: An Induced Fit Docking Approach for Full Receptor Flexibility. J Chem Inf Model. 2016 Apr 25;56(4):734–46.

75. Atilgan C, Gerek ZN, Ozkan SB, Atilgan AR. Manipulation of conformational change in proteins by single-residue perturbations. Biophys J. 2010 Aug 4;99(3):933–43.

76. Li Z, Bolia A, Maxwell JD, Bobkov AA, Ghirlanda G, Ozkan SB, Margulis CJ. A Rigid Hinge Region Is Necessary for High-Affinity Binding of Dimannose to Cyanovirin and Associated Constructs. Biochemistry (Mosc). 2015 Nov 24;54(46):6951–60.

77. Day WHE, Edelsbrunner H. Efficient algorithms for agglomerative hierarchical clustering methods. J Classif. 1984 Dec 1;1(1):7–24.

